# The nucleolar aberrancies that drive ribosome impairment induced by RNA binding proteins are hallmarks of aging

**DOI:** 10.1101/2023.03.07.531270

**Authors:** Pedro Aguilar-Garrido, María Velasco-Estévez, Miguel Ángel Navarro-Aguadero, María Hernandez-Sanchez, Prerna Malaney, Xiaroui Zhang, Marisa J. L. Aitken, Alvaro Otero-Sobrino, Marta Ibañez-Navarro, Alejandra Ortiz-Ruiz, Diego Megias, Manuel Pérez-Martínez, Gadea Mata, Jesús Gomez, Orlando Dominguez, Osvaldo Graña-Castro, Eduardo Caleiras, Paloma Jimena de Andres, Sandra Rodriguez, Raúl Torres, Oleksandra Sirozh, Vanesa Lafarga, Joaquín Martinez-Lopez, Sean M. Post, Miguel Gallardo

## Abstract

The nucleolus is a dynamic structure where ribosome subunits are produced. Indeed, nucleoli respond to any change in cellular homeostasis by altering the rate of ribosome biogenesis, thus working as a stress sensor. Therefore, an imbalance in ribosome biogenesis promotes changes in morphology and function and can evoke a nucleolar stress response. Changes in the structure and composition of nucleoli impair ribosome biogenesis and have been described as nucleolar stress, a mechanism related to aging and cancer.

Here, we show the role of the RNA binding protein Hnrnpk in nucleolar dynamics and ribosome function. Hnrnpk is a ribonucleoprotein in charge of escorting nascent transcripts to its processing and nuclear export to ribosomes. When Hnrnpk is overexpressed, the nucleolus is altered and shows stress-like phenotype, with accumulation and delocalization of components such as Ncl, driving ribosome biogenesis impairment and halting protein translation.

Nucleolin haploinsufficiency is correlated with enlarged nucleoli, increased ribosome components and translation and induces a reduction in lifespan. Thus, gain of Ncl generated by Hnrnpk overexpression can cause ribosome biogenesis defects associated with ribosome impairment leading to ribosomopathies and bone marrow failure syndrome.

Aging and bone marrow failure share common biological hallmarks. Indeed, Hnrnpk overexpression and nucleolar stress trigger cell cycle arrest and senescence of the cells, a feature of both processes.

Together, these findings support the idea that nucleolar abnormalities contribute to ribosome impairment, thus triggering the onset of hematopoiesis and the aging process. Here, we decipher a novel master regulator of this mechanism: Hnrnpk.

## Introduction

The nucleolus is the biological structure where ribosome subunits are generated by the transcription of rDNA, rRNA processing and ribosome subunit assembly. The nucleolus is divided into three structures: the fibrillary centers (FC), the dense fibrillary component (DFC) and the granular component (GC). rDNA transcription occurs in FC and the boundaries of FC and DFC, while assembly of rRNA and ribosome proteins of 40S and 60S ribosome subunits are performed in the GC, which will be later exported to the cytoplasm for their final assembly into a functional ribosome. Thus, ribosome biogenesis and nucleoli are interconnected as a feedback loop. Indeed, aberrancies in nucleolus function and composition alter ribosome function and protein translation, as dysregulation of ribosome biology impacts nucleolus organization. Therefore, ribosome stress (RS) and nucleolar stress (NS) are dependent and complementary responses^1,2^.

The rRNAs transcribed and processed in the nucleoli make up to 80% percent of the total RNA generated on the cell, carrying an enormous energetic cost. Hence, nucleolar activity is associated with caloric consumption, metabolism and ribosome production. Nucleolus plays a role in lifespan regulation and it has been observed that a reduction in some of its components, such as fibrillarin, is correlated with a prolonged lifespan^3^. This finding is in line with the fact that a reduction in ribosome biogenesis regulatory molecules, such as mTOR or c-MYC, also promotes the lifespan of organisms^4,5^. Thus, as a central paradigm, when the ribosome components are well balanced, smaller nucleoli and a reduction in ribosome biogenesis and translation are associated with prolonged lifespan; while enlarged nucleoli, an increase in ribosome number and elevated protein translation are associated with cancer and aging processes, with similar phenotypes observed when ribosome components are unbalanced^2^.

RNA-binding proteins (RBPs) are a diverse class of molecules that interact with transcripts and noncoding RNAs^6^. They are commonly associated with ribonucleoprotein complexes (RNPs) and bind nascent transcripts for processing and nuclear export to ribosomes^7^. RBPs are involved in each step of the post-transcriptional processing of transcripts, dictating their fate and function and, through RNPs, regulating a plethora of biological functions such as splicing, polyadenylation, stability localization, translation and degradation^8^. These proteins can be found in different biological structures, such as the nucleoli where the RBP Ncl has a critical role in nucleolus activity, rRNA synthesis and cell lifespan^3,9^. However, RBPs usually shift between compartments, as occurs for Ncl and Hnrnpk, a poly(C)-binding ribonucleoprotein that tenaciously binds C-rich tracks of DNA and RNA, both of which fluctuate between the nucleus and cytoplasm to export nascent transcripts^10-12^. Thus, RBPs are key participants in nucleolus/ribosome cross-talk. Indeed, multiple RBPs are associated with nucleolar biology^13^ and have been suggested to exert a role in aging^14,15^.

Hematopoietic stem cells (HSCs) are especially sensitive to the aging process. Aged HSCs lose their amplification and proliferation capacity, promoting their exhaustion and final differentiation^16^. There are multiple causes of HSC exhaustion, one of which is ribosome dysregulation that occurs in ribosomopathies. These abnormalities are consistent with different bone marrow failures such as myelodisplastic syndrome (MDS), Diamond–Blackfan anemia (DBS) or Shwachman-Diamond Bodian Syndrome (SDS). Genetically, they are frequently associated with deleterious mutations in RBPs (e.g EMG1, UTP4) ribosomal proteins (e.g. RPS14, RPSA) or rRNAs interacting proteins (SBDS, eIFs)^17^. Therefore, RBPs are usually found to be altered in HSC biology, affecting ribosome biogenesis and function.

Here we show, without precedent, that the overexpression of RBPs, such as Hnrnpk, can also cause nucleoli abnormalities by driving ribosome impairment in both CRISPR-generated cell models and murine models. This ribosome dysfunction reduces mouse lifespan *in vivo* by driving bone marrow failure syndrome similarly to what is observed in ribosomopathies. Therefore, this study reveals novel connections between nucleoli, RBPs and ribosomes; lifespan and ribosomopathies; and how these connections all have an impact on aging, especially in the hematological compartment.

## Results

### The Hnrnpk-overexpressing mouse model has lower tumor onset and a lifespan reduction *in vivo*

To study the impact of Hnrnpk overexpression in an *in vivo* context, we developed the *Hnrnpk*^Tg^ mouse. To avoid developmental issues reported by Hnrnpk, we used the inducible tamoxifen (TAM) UBC^creERT2^ model which was activated 3-6 weeks after birth (Figure 1A). The activated *Hnrnpk*^Tg/Ubc-creERT2^ mouse model showed a detectable overexpression of hnRNP K protein in cells and tissues such as bone marrow, spleen or liver, as determined by IHC and/or western blot (Figure 1B). Consistently, the *Hnrnpk*^Tg/Ubc-creERT2^ model had a reduction in lifespan (Figure 1C), mainly due to dysplasia and/or bone marrow failure (Figure 1D).

**Fig. 1.**
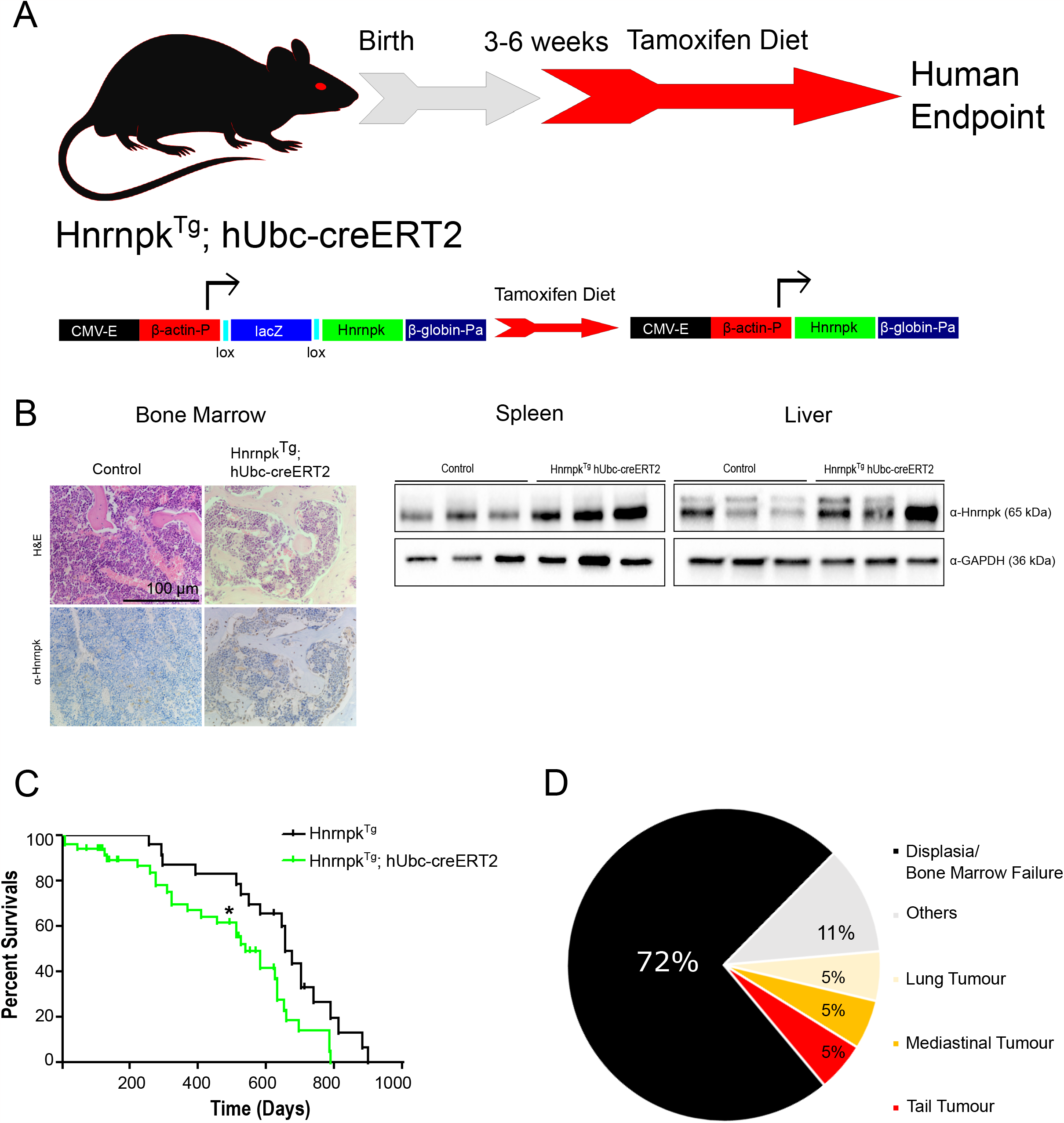
Hnrnpk^Tg/Ubc-creERT2^ mouse model phenotype. (A) Flow chart of *Hnrnpk*^Tg^ and *Hnrnpk*^Tg/Ubc-creERT2^ mouse models. (B) Bone marrow IHC from *Hnrnpk*^Tg/Ubc-creERT2^ mice and western blot membrane showing Hnrnpk overexpression in *Hnrnpk*^Tg/Ubc-creERT2^ liver and spleen tissues (n=3). (C): Kaplan–Meier survival curve of *Hnrnpk*^Tg/Ubc-creERT2^ mice (TAM ON) vs *Hnrnpk*^Tg^ (TAM ON) (*Hnrnpk*^Tg/Ubc-creERT2^ =49 vs *Hnrnpk*^Tg^=22, p=0.032 HR:0.5305). (D) Pie-chart with cause of death/phenotype developed for *Hnrnpk*^Tg/Ubc-creERT2^ mice (n=14)

### The Hnrnpk mouse model develops bone marrow failure and an aging phenotype

During the phenotypical characterization of our novel *Hnrnpk*^Tg/Ubc-creERT2^ mouse model, we identified different signs of aging: a higher percentage of *Hnrnpk*^Tg/Ubc-creERT2^ mice suffered skeletal deformities and kyphosis and white or bald hair patches (Figure 2A).

**Fig. 2.**
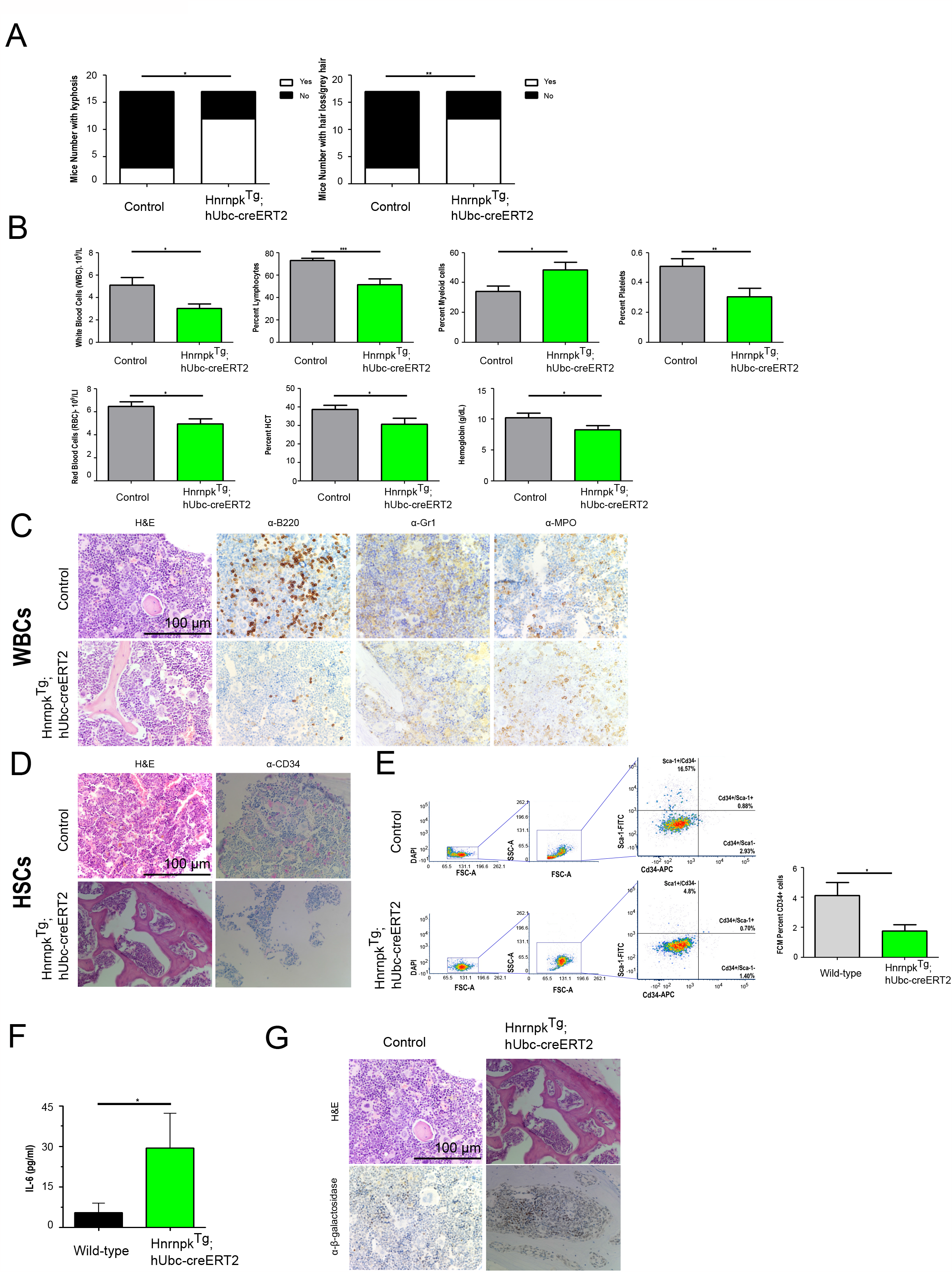
hnRNP K overexpression drives bone marrow failure and aging in vivo. (A) Aging characteristics by box-plots % showing incidence of skeletal dephormities/kyphosis (p=0.011) and white/bald hair patches (p=0.001) (n=14 vs 9, Fisher analysis). (B) CBC of total WBCs (p=0.032), % lymphocytes (p=0.001), % myeloid cells (p=0.017) and % platelets (p=0.009), total red blood cells (p=0.016), hematocrit (p=0.050) and hemoglobin (p=0.017) (n=14 vs 9, two-sided Student’s t-test analysis). (C) Mature white blood cell IHC analysis: H&E and B220, Gr-1 and MPO. Scale bar: 100 μm. (D) HSCs analysis: H&E and CD34. Scale bar: 100 μm. (E) FCM gating strategy, dot plot Cd34/Sca-1 analysis of total bone marrow from TAM age-paired 300 days and box plot showing percent Cd34 positive cells (*Hnrnpk*^Tg/Ubc-creERT2^ =3 vs *Hnrnpk*^Tg^=4, p=0.046) (F) Senescence analysis: ELISA Il-6 box-plot concentration (TAM age-paired *Hnrnpk*^Tg/Ubc-creERT2^ =9 vs *Hnrnpk*^Tg^=5, p=0.046) and (G) IHC β-galactosidase staining in bone marrow. All graphs are shown as mean ± SD. All images are representative of at least n=3 mice and 4 random pathological areas.

Additionally, we characterized the hematological compartment of the mice. By complete blood count (CBC) analysis, we observed the development of leukopenia, lymphopenia, anemia and thrombocytopenia in *Hnrnpk*^Tg/Ubc-creERT2^ mice (Figure 2B). We then analyzed the hematological tissues bone marrow and spleen. H&E/IHC and FCM analysis of the bone marrow showed several abnormalities, including a reduction in lymphoid B220+ cells, a slight increase in myeloid lineage Gr1+ and Mpo+ cells and improper hematopoiesis, with a reduction in CD34+ and Sca1+ cells (Figure 2C-E). Furthermore, we observed an increase in the expression of the senescence marker β-galactosidase, as well as higher levels of the pro-senescent cytokine Il-6, as measured by ELISA (Figure 2F-G).

### Hnrnpk overexpression promotes cell cycle arrest and senescence

To understand the molecular impact of hnRNP K overexpression that drives the bone marrow failure phenotype *in vivo*, we developed an Hnrnpk overexpressing *in vitro* model using mouse embryonary fibroblasts (MEFs) and CRISPR-Cas9 synergistic activator mediator (SAM) technology, obtaining MEFs that constitutively overexpressed Hnrnpk (Figure 3A-B). To continue, we developed an RNA-seq analysis of Hnrnpk overexpressing MEFs. The most significant differential expression was observed in the Molecular Signatures Database (MSigDB)^18^ G2/M checkpoint pathway, with an upregulation of pathway genes in the Hnrnpk overexpression model (Figure 3C). Next, to determine aberrancies in the G2/M checkpoint pathway, we conducted a cell cycle FCM study, confirming an increase in G2/M phase and polyploid cells in our Hnrnpk overexpression model (Figure 3D-F). To determine whether this cell cycle arrest drove cells to senescence, we developed an SA-β-galactosidase analysis, confirming an increase in senescence (Figure 3G-H). Finally, p53/p21, -which is directly regulated by hnRNP K- and p16, all of which are key molecules in the regulation of cell cycle arrest, were elevated in hnRNP K overexpressing cells (Figure 3I). Taken together, these results suggest that hnRNP K promotes cell cycle arrest, driving the cell to a senescent state.

**Figure 3.**
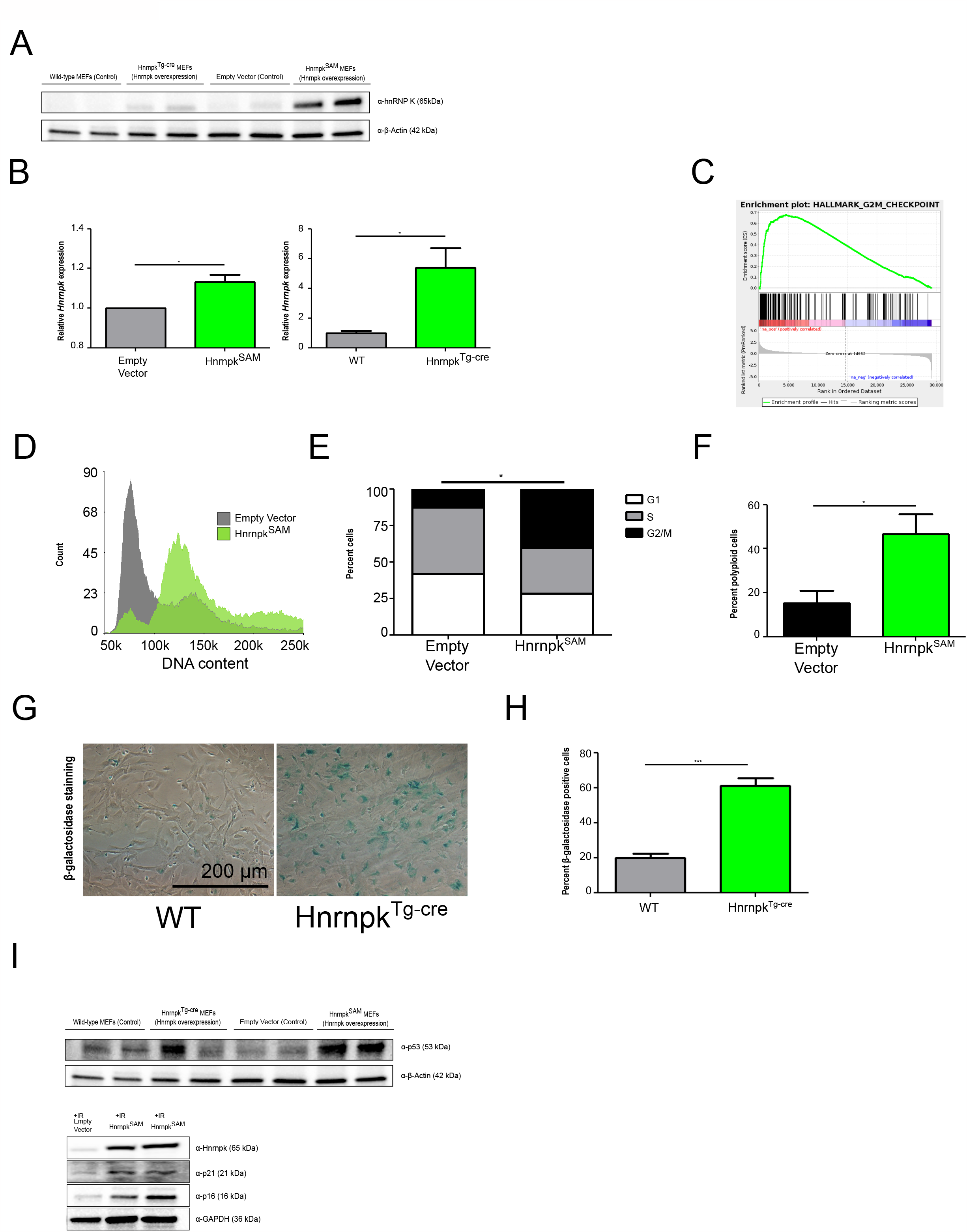
Hnrnpk drives senescence. (A) Western blot membrane showing Hnrnpk overexpression in *Hnrnpk*^*SAM*^ and *Hnrnpk*^*Tg-cre*^ MEFs. (B) qRT-PCR results box-plot showed Hnrnpk overexpression in *Hnrnpk*^*SAM*^ MEFs (p=0.05) and *Hnrnpk*^*Tg-cre*^ (p=0.014). (C) Gene Set Enrichment Analysis (GSEA) enrichment plot of the top 1 significantly regulated pathway (MsigDB): HALLMARK_G2M_CHECKPOINT (FDR-qval<10^−4^) in transcriptomes of *Hnrnpk*^SAM^ vs Empty vector MEFs (n=3). A false discovery rate q-value of <0.25 was chosen as a cut-off for exploratory data analysis (D) Hnrnpk overexpressing cells showed an increase in the G2/M and polyploidy populations measured by cell cycle FCM plots (E) Box-plot analysis of G1, S and G2/M % cells from FCM analysis (p=0.01) (F) and polyploidy (p=0.02). (G) Bright-field microscope images of SA-β-galactosidase staining in HnrnpK overexpressing MEFs. (H) Box-plot of positive cells for SA-β-galactosidase staining (p=0.0001). (I) Western blot membrane of Hnrnpk overexpressing cells showed higher p53 expression (top). Western blot membrane of Hnrnpk overexpressing cells with irradiation showed an increase in senescence markers p21 and p16 (bottom). All graphs are shown as mean ± SD. All statistical analyses were two-sided Student’s t-test, with the exception of FCM cell cycle analysis (two-way ANOVA). All experiments comprised at least n=3 biological replicates and/or n=3 technical replicates.

### Hnrnpk overexpression drives ribosome dysfunction

Once we determined the senescent phenotype of Hnrnpk-overexpressing cells, we aimed to decipher the molecular mechanisms that trigger cell cycle arrest. Interestingly, the RNA-seq analysis of our Hnrnpk overexpression model showed an enrichment of gene sets related to ribosome biogenesis, such as mTORC1 and Myc (Enrichr graph-bars)^19^, and the significance of Molecular Signatures Database (MSigDB)^18^ mTOR (Figure 4A). Thus, Hnrnpk expression seemed to directly correlate with ribosome biogenesis master regulators c-Myc and mTor (Figure 4B) and downstream activation of targets of mTor, such as 4Eb-p1.

**Fig. 4.**
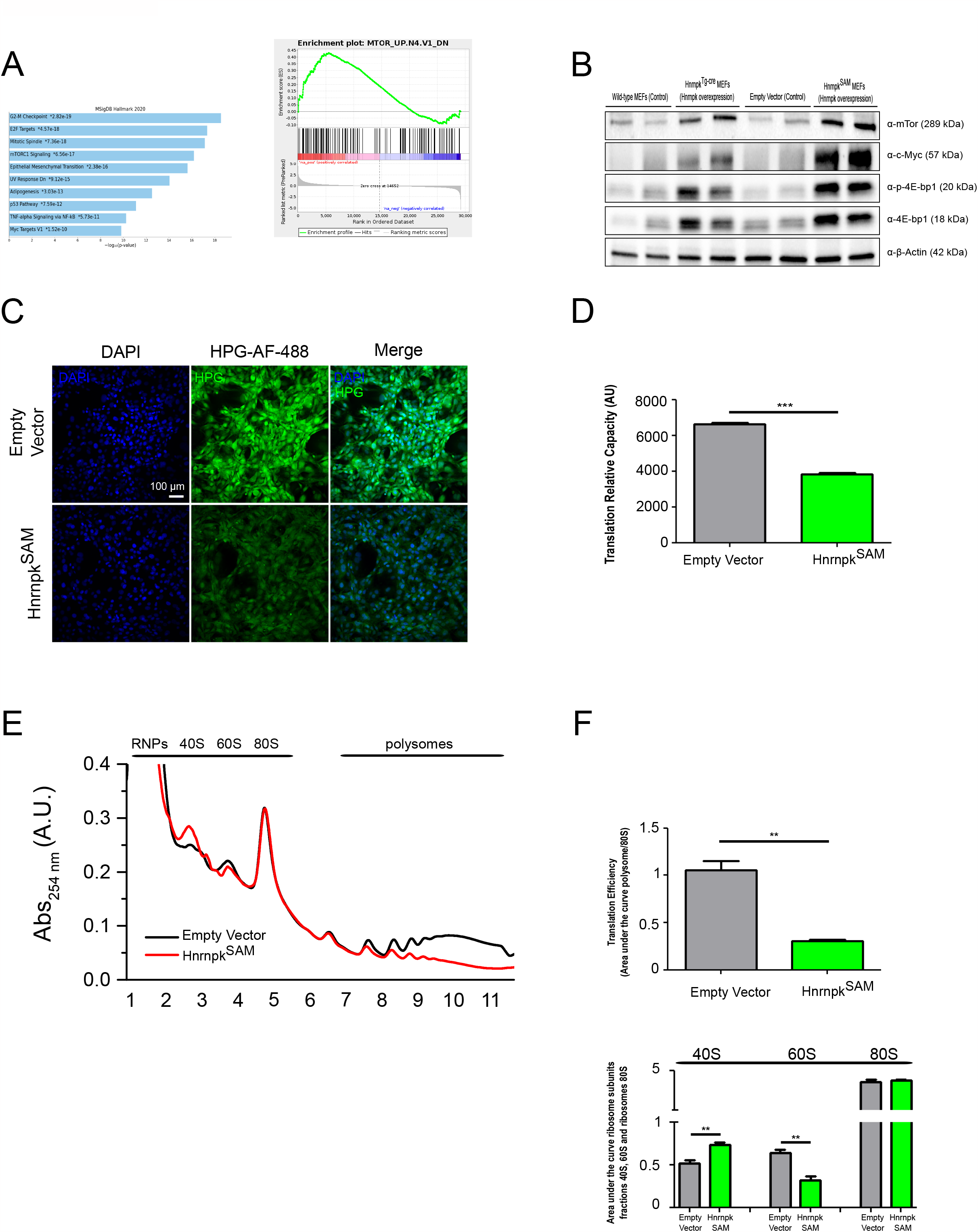
hnRNP K regulates ribosome biogenesis and function. (A) Enrichr Molecular signature database hallmarks bar-graph (left) and Gene Set Enrichment Analysis (GSEA) enrichment plot of significantly regulated pathways (MsigDB): MTOR_UP.N4.V1_DN (FDR-qval<10^−4^) from RNA-seq data significant differentially expressed genes (FDR < 0.05) (B) Western blot membrane showing that Hnrnpk overexpression directly correlates with c-Myc and mTor expression, and mTor downstream target 4Eb-p1 and its activated form (p-Eb-p1) showed a increase in Hnrnpk overexpression MEFs (p=0.0079). (C) HPG assay confocal microscopy from hnRNP K overexpressing *Hnrnpk*^SAM^ MEFs (D) Fluorescence intensity values from the HPG assay showed decreased global translation in Hnrnpk-overexpressing MEFs (p=0.0007). (E) Polysome assay profiles shows a decrease in polysome fraction, and unbalance in ribosome subunits (40S p=0.008; 60S p=0.005). (F) Translation efficiency values (area under the curve of polysome fraction/area under the curve of 80S fraction) shows a decrease in Hnrnpk overexpression MEFs (p=0.002). Scale bar: 100 μm. All graphs are shown as mean ± SD. PCR statistical analysis was two-sided Student’s t-test. All studies comprised at least n=3 biological replicates and/or n=3 technical replicates.

These results suggest that Hnrnpk overexpression can cause an increase in ribosome biogenesis and ribosome functions. To confirm this hypothesis, we evaluated ribosome function by global protein synthesis through an HPG assay (Click-iT™ HPG assay; ThermoFisher, #C10428). Surprisingly, our results pinpointed a halt in global translation in Hnrnpk-overexpressing cells (Figure 4C-D). Ribosome biogenesis dysfunction is commonly associated with ribosomopathies, abnormalities that course with bone marrow failure and drive ribosomal function defects and senescence phenotypes.

Therefore, we next evaluated the translation efficiency and ribosome subunit components with a polysome fractionation assay. We observed that Hnrnpk-overexpressing cells had a decrease in translation efficiency (Figure 4E-F). This result confirmed not only a reduction in ribosome biogenesis with a lower quantity of active ribosomes, but also an increase in inactive ribosomes and/or ribosome subunits, thus causing the ribosomopathy phenotype.

### Hnrnpk overexpression drives nucleolar abnormalities with a nucleolar stress signature

The combination of a deficiency in translation and ribosome biosynthesis with an overexpression of ribosome biogenesis master regulators such as mTor and c-Myc, suggested the existence of a ribosomopathy. Thus, we hypothesized that ribosome component synthesis, processing or assembly could be compromised. Hence, we focused on nucleolus, the structure with the mission of rRNA synthesis and ribosome component assembly. Indeed, our RNA-seq results showed an enrichment of genes belonging to the nucleus and nucleolus (*data not shown*).

The first impressions obtained from Hnrnpk-overexpressing cell images were an enlargement of nucleous and DNA quantity, consistent with the polyploidy and G2/M arrested senescent phenotype observed by FCM (Figure 5A-B). We then studied the nucleoli. The nucleolus is a non-layer biological structure with different components and a myriad of proteins involved. Hnrnpk-overexpressing cells showed nucleolar abnormalities in some of these components, such as Ncl, a protein found in DFC and GC that is involved in rRNA synthesis, processing and ribosome assembly. We observed that Ncl was overexpressed and delocalized onto the nucleoplasm (Figure 5A-D).

**Fig. 5.**
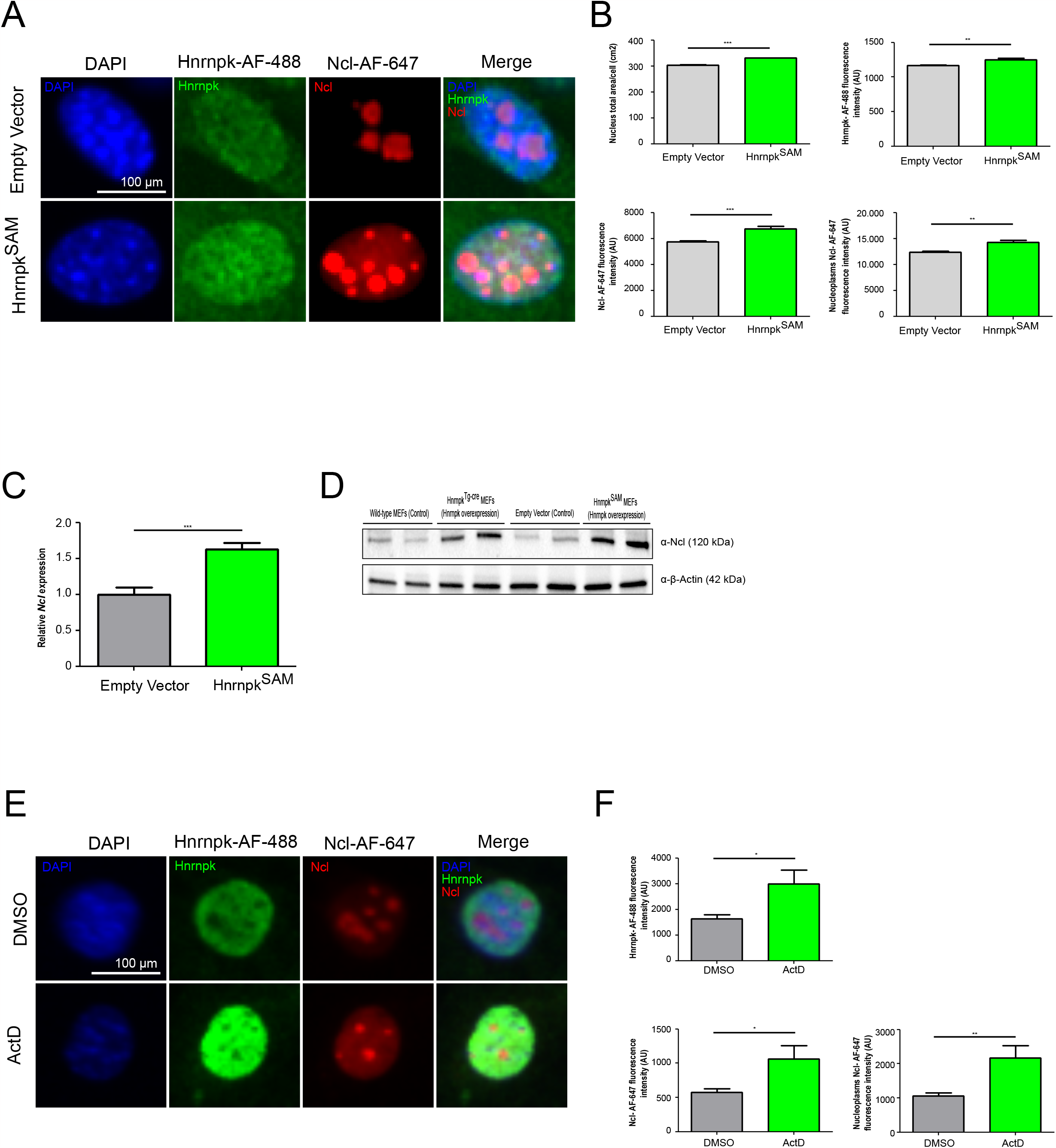
hnRNP K overexpression dysregulates nucleolar components and triggers nucleolar stress signs. (A) Images of confocal microscopy with Ncl, Hnrnpk and DAPI staining in wild-type and *Hnrnpk*^*Tg-cre*^ MEFs. (B) Box-plot analysis of: total area (DAPI, nucleus, p=0.0001), Alexa Fluor 488 intensity (Hnrnpk, p=0.0016) and Alexa Fluor 647 intensity (Ncl, p=0.0009); and nucleoplasm Alexa Fluor 647 intensity (Ncl delocalization, p=0.0025). (C) qRT-PCR results box-plot showing Ncl overexpression in *Hnrnpk*^*Tg-cre*^ MEFs (p=0.0005). (D) Western blot membrane showing that Hnrnpk overexpression directly correlates with Ncl overexpression. (E) Confocal microscopy images of Ncl, Hnrnpk and DAPI staining in wild-type and hnRNP K-overexpressing cells treated with ActD (5nM). (F) Top: Box-plot analysis of Alexa Fluor 488 intensity (Hnrnpk) and Alexa Fluor 647 intensity (Ncl) Bottom: Nucleoplasm Alexa Fluor 647 intensity (Ncl delocalization) in MEFs treated with ActD (5 nM) (Hnrnpk: p=0.0381; Ncl: p=0.0190; Ncl delocalization: 0.0095). Scale bar: 100 μm. All graphs are shown as mean ± SD. Statistical analysis consisted of two-sided Student’s t-test. All experiments comprised at least 2 biological replicates and/or 2 technical replicates.

Nucleolin is a recognized stress sensor and an increase in Ncl is frequently correlated with nucleolar stress. To confirm that Hnrnpk overexpression shows nucleolar stress hallmarks, we induced nucleolar stress in wild-type MEFs by actinomycin D treatment (ActD). Nucleolar stress induction promoted Hnrnpk overexpression and mimicked this signature: an increase in Ncl expression and delocalization onto the nucleoplasm (Figure 5E-F).

Collectively, our data indicated that Hnrnpk overexpression drives nucleolar abnormalities with a signature similar to that of the nucleolar stress response, driving ribosome dysfunction and a decrease in translation efficiency, common hallmarks of ribosomopathies. Stress signals trigger the p53/p21 pathway, leading to cell cycle arrest and promoting senescence. The consequence *in vivo* is the typical signature of a ribosomopathy, with a bone marrow failure phenotype and hallmarks of aging.

## Discussion

Altogether, our work establishes Hnrnpk as a regulator of nucleolar/ribosome crosstalk, where upregulation of this RBP strikingly promotes nucleolar abnormalities with an impact on ribosome biogenesis and consequent ribosome dysfunction, inducing aging and disrupting regular hematopoiesis.

Hnrnpk overexpression induces abnormalities in the nucleolus and these cells share common hallmarks of nucleolar stress such as the delocalization and imbalance in nucleolar proteins. In this regard, we found that nucleolin (Ncl), a DFC and GC nucleolar protein related to nucleoli activity and lifespan^3,9^, was overexpressed and delocalized in Hnrnpk-overexpressing cells. Indeed, we had previously determined the direct interaction between Hnrnpk and Ncl proteins, suggesting a straight regulation^20^. Therefore, we and others ascertain that Hnrnpk regulates the protein, mRNA or DNA levels of both c-Myc and mTor^20,21^. Both molecules are conspicuously linked to the regulation of rRNA and ribosome protein synthesis, processing and assembly^3^ and a reduction in mTor and c-Myc levels drives an increase in lifespan^4,5^.

Here, we propose Hnrnpk as a pivotal candidate in ribosome biogenesis and function. Hnrnpk positively regulates Ncl. Ncl haploinsufficiency is correlated with enlarged nucleoli, an increase in ribosome components and translation, and a reduced lifespan^3^. However, in our model, Ncl overexpression is correlated with the opposite mechanism, a decrease in ribosome components and translation, but with the same consequence, a reduction in lifespan. This evidence supports the idea that Ncl is a master regulator of ribosome biology and that any imbalance drives nucleolar/ribosome abnormalities and the aging phenotype. Indeed, Ncl is involved in a myriad of rRNA processes, including the regulation of rDNA transcription by negative regulation of RNA POLI, but also processing or transport^22^. Thus, Ncl overexpression induced by Hnrnpk could negatively impact rRNA, and, consequently, ribosome component production promoting an imbalance and generating “orphan” ribosomal components that are not assembled into a fully active ribosome, thus generating a ribosomopathy phenotype.

Numerous studies have revealed the link between ribosome biogenesis reduction and increased longevity through dietary restriction and reduction of energetic cost^2,23,24^. However, defective ribosome biogenesis also leads to ribosomopathies and a dramatic reduction in lifespan by affecting specific tissues such as the hematopoietic compartment in an aging-like fashion. Furthermore, a reduction in rDNA copy number and/or rRNAs (which occurs in some ribosomopathies) leads to overexpression of mTor, suggesting a feedback loop between them^25^. Indeed, other ribosome biogenesis master regulators, such as c-Myc, regulate RNA POLI. Thus, it is logical to think that, when the reduction in RNA POLI activity caused by the Hnrnpk-Ncl overexpression loop occurs, the cell will try to compensate for this lack of activity and loss of rRNA synthesis with an overexpression of molecules such as mTOR and c-Myc. This overexpression of mTOR and c-Myc could contribute to lifespan shortening, the opposite effect that occurs when these molecules are haploinsufficient and prolong lifespan. Therefore, we have described here that the upregulation of Hnrnpk positively regulates Ncl, and their overexpression leads to ribosome biogenesis imbalance and translation deficiency, which, in a compensatory attempt, upregulates mTOR and c-Myc molecules, leading to lifespan shortening.

The nucleolus is also the site for the assembly of ribonucleoprotein particles such as Hnrnpk, thus regulating numerous functions such as splicing, telomere biology, stress granule biogenesis and long/small noncoding RNA machinery, driving genome integrity, stress signaling or the cell cycle^26^. Indeed, Hnrnpk is a key player in the regulation of DNA damage, a stress signal^27-29^. In this work, we showed a connection between nucleolar abnormalities, ribosome dysfunction and cell cycle arrest through the p53/p21 pathway, thus, leading to a senescent phenotype, similar to what has been observed with c-Myc or mTor overexpression in other works^30,31^.

Aging is a complex biological process with multiple mechanisms involved. There is a plethora of data revealing that aged cells suffer cell cycle arrest and have a senescence fingerprint (e.g. SA-β-galactosidase positive, p21 overexpression and pumped Il-6 levels)^32^. Hence, the senescence phenotype driven by Hnrnpk overexpression could be the cause of lifespan reduction in our cellular and animal model.

One of the most frequent signs of aging is inefficient hematopoiesis, with a reduction in hematopoietic stem cells which triggers anemia and cytopenia. There are different hematological syndromes that course with bone marrow failures, as observed in myelodysplastic syndrome (MDS), Blackfan-Diamond Anemia (BDA) or Shwachman-Diamond Bodian Syndrome (SDS). Molecularly, harmful loss-of-function mutations in ribosome proteins, biogenesis factors, rRNA or rDNA transcription defects are frequently linked to ribosomopathies, in which aberrancies in rRNAs and/or ribosomal proteins lead to incorrect ribosome biogenesis, as the subunits cannot be assembled properly^17^. Here, we showed a novel mechanism that triggers ribosomopathies: a gain of expression of RBPs that leads to aberrancies and unbalancing of rRNAs and/or ribosome proteins, consequent ribosome dysfunction and a bone marrow failure phenotype. Indeed, our animal model mimics the anemia, thrombocytopenia and lymphopenia usually observed in bone marrow failure but also presents other signs of ribosomopathies, such as the skeletal disorders found in SDS. Moreover, bone marrow failure is also characterized by increased cell cycle arrest and senescent cells, as well as IL-6 cytokine release, which can also abrogate lymphoid lineage and promote myeloid differentiation^33,34^ similar to what was observed in our model.

Our work identifies a link between nucleolar abnormalities and longevity. Regulation of ribosome biogenesis and function are crucial elements that are regulated by the nucleolus. RNA-binding proteins and ribonucleoproteins such as Hnrnpk and Ncl are key players in rRNA and RP synthesis and processing and subsequently, key players in nucleolar/ribosome crosstalk. These processes implicated in longevity could be similarly applied to hematopoietic stem cell biology, which are particularly sensitive to ribosome dysregulation as it occurs in ribosomopathies and in our novel ribosomopathy model. Conversations between the ribosome and the nucleoli coordinate cell cycle progression^1,2,35^. Here, we prove that nucleolar abnormalities and ribosome dysfunction can induce cell cycle arrest and poliploidy driving towards a senescence phenotype that can explain the reduction in lifespan and bone marrow failure found in our novel mouse model.

In conclusion, our work shows, without precedent, that overexpression of RBPs, such as Hnrnpk or Ncl, triggers ribosomopathies. This is in contrast with the current accepted paradigm that establishes that loss of ribosomal molecules causes these syndromes. This evidence highlights that the subyacent cause of ribosomopathy is the imbalance and dysregulation of ribosome biology in any direction, either defective or upregulated. Thus, aging cells and ribosomopathies would benefit from precision medicine approaches, particularly, from the therapeutic exploitation of Hnrnpk, the inhibition of nucleolar-related molecules regulated by Hnrnpk such as Ncl (aptamer AS1411)^36^ and, potentially, ribosome biogenesis regulators such as c-Myc (i.e., Omomyc, bromodomain inhibitors)^20,37^ or mTor (i.e., everolimus)^38^.

## Methods

### hnRNP K-overexpressing animal model generation

hnRNP K^Tg^ was generated at MD Anderson Cancer Center (MDACC) by Dr. Sean M. Post.

To generate a conditional Hnrnpk overexpression model, we cloned full-length Hnrnpk as a BglII-XhoI fragment into the pCALL2 vector. This construct contains a chicken β-actin promoter with an upstream cytomegalovirus (CMV) enhancer (pCAGGS). This promoter is followed by a loxP-flanked LacZ/neoR fusion with three SV40 polyadenylation (PA) signals and the full-length Hnrnpk cDNA. We refer to the expression vector as pCALL2-Hnrnpk. The linearized vector was injected into blastocysts to generate chimeric mice by the Genetically Engineered Mouse Facility at the MD Anderson Cancer Center. Transgenic mice were mated with C57Bl/6 mice to produce transgenic offspring. The pCALL2-Hnrnpk transgene was genotyped by real-time PCR on DNA extracted from tail clips.

The generation of the mouse model was conducted in the MD Anderson Genetically Engineered Mouse Facility (GEMF) with approval from the Institutional Animal Care and Use Committee at MD Anderson Cancer Center under protocol 0000787-RN01.

Hnrnpk^Tg^ mice were crossed with a mouse strain carrying ubiquitously expressed, tamoxifen-activated recombinase, hUBC-CreERT2, to generate Hnrnpk^Tg^ and Hnrnpk^Tg-hUBC-creERT2^ mice. These mice were fed *ad libitum* with a tamoxifen-containing diet for the long term, starting at 3-6 weeks of age. All mice were maintained at the Spanish National Cancer Centre under specific pathogen-free conditions in accordance with the recommendations of the Federation of European Laboratory Animal Science Associations (FELASA). All animal experiments were approved by the Ethical Committee (CEIyBA) from the Spanish National Cancer Centre and performed in accordance with the guidelines stated in the International Guiding Principles for Biomedical Research Involving Animals, developed by the Council for International Organizations of Medical Sciences (CIOMS), under the protocol PROEX158/18.

### Mice genotyping and validation of the *Hnrnpk* allele

PCR-based strategies using primer sets that were external and internal to both the 5′ and 3′ arms were initially performed to confirm homologous recombination and germline transmission. The 5′ arm was verified by PCR amplification and visualization using external and internal primers.

The pCALL2-Hnrnpk transgene genotyping was performed by real-time PCR allelic discrimination on genomic DNA at the CNIO GMM Genotyping service with the following primers and hydrolysis probe: Hnrnpk: F_30F12 CCAGATACAGAACGCACAGT, pCALL:R_30F13 AAGGGGCTTCATGATGTCC and pCALL:S_30F14 Fam-CTCGAGGTGGCTGCGATC-Zen-IBFQ.

### Pathological Analysis and Immunohistochemistry

Tissue samples were fixed in 10% neutral buffered formalin (4% PFA in solution), paraffin-embedded and cut at 3 μm, mounted in TOMO slides and dried overnight. For different staining methods, slides were deparaffinized in xylene and rehydrated through a series of graded ethanol until water. Consecutive sections were stained with hematoxylin and eosin (H&E), and several immunohistochemistry reactions were performed in an automated immunostaining platform (Autostainer Link 48, Dako; Ventana Discovery XT, Roche).

Antigen retrieval was first performed with the appropriate pH buffer, (low or high pH buffer, Dako; CC1m, Ventana, Roche) and endogenous peroxidase was blocked (peroxide hydrogen at 3%). Then, slides were incubated with the appropriate primary antibody as detailed in Table 1.

**Table 1:** IHC antibodies

| Host | Name | Clone | Dilution | Company | Catalogue |
| --- | --- | --- | --- | --- | --- |
| Mouse Monoclonal | Hnrnpk | D-6 | 1/400 | Santa Cruz Biotechnology | Sc-28380 |
| Rat Monoclonal | Cd34 | RAM34 | 1/100 | Invitrogen | 14-0341-81 |
| Rabbit Monoclonal | c-kit | EPR22566-205 | 1/200 | Abcam | ab231780 |
| Rabbit Polyclonal | Mpo |  | 1/1500 | Dako | A0398 |
| Rat Monoclonal | Gr1 | RB6-8C5 | 1/100 | Invitrogen | 14-5931-82 |
| Rat Monoclonal | Cd11b | M1/70 | 1/1000 | eBioscience | 14-0112-82 |
| Rat Monoclonal | CD45R/B220 | RA3-6B2 | 1/100 | Becton Dickinson co. | 01121A |
| Rat Monoclonal | p21 | 291H | 1/10 | Abcam | Ab107099 |
| Rat Monoclonal | B-galactosidase | 3A9A10F8 | 1/1000 | Monoclonal antibody core unit | CNIO |

After primary antibody incubation, slides were washed and incubated with the corresponding secondary antibodies when needed (rabbit anti mouse, Abcam; rabbit anti rat, Vector; Goat anti-rabbit, Dako) and visualization systems (Bond Polymer Refine Detection, Bond, Leica; Omni Map anti-(rabbit,mouse,rat) Ventana, Roche) conjugated with horseradish peroxidase.

Immunohistochemical reactions were developed using 3, 30-diaminobenzidine tetrahydrochloride (DAB) (Chromo Map DAB, Ventana, Roche; DAB Dako) and nuclei were counterstained with Carazzi’s hematoxylin. Finally, the slides were dehydrated, cleared and mounted with permanent mounting medium for microscopic evaluation.

Positive control sections known to be primary antibody positive were included for each staining run.

### Il-6 ELISA quantification

Peripheral blood of euthanized mice was treated with BD Pharm LyseTM buffer (BD Biosciences, #555899) and centrifugued, and serum was frozen at -80°C until measurement. Enzyme-linked immunosorbent assay (ELISA) was used to determine the levels of secreted Il-6 in mouse blood plasma (ELISA) using a Mouse Il-6 DuoSet ELISA Kit (R&D, DY406). Measurement and analysis of Il-6 levels were carried out according to the manufacturers’ instructions.

### Generation of hnRNP K overexpressing mouse embryo fibroblasts (mouse-derived and CRISPR/SAM-edited)

Mouse embryo fibroblasts (MEFs) from WT and Hnrnpk^Tg-cre^ (derived from the breed of Hnrnpk^Tg^ with Hnrnpk^Ella-cre^ or Hnrnpk^Ubc-creERT2^) were prepared from embryonic day 13.5 embryos and cultured in Dulbecco’s modified Eagle’s medium supplemented with 10% fetal calf serum at 37°C and 5% CO2. Additionally, to generate hnRNP K-overexpressing cellular models, wild-type MEFs were transduced with lentiviral particles containing the plasmids lentiMPHv2 (Addgene #89308) and lentiSAMv2 (Addgene #75112). At 72 hr post-transduction, the cells were selected by blasticidin (2 μg/mL) (AG Scientific, CA, USA, Cat# B-1247) and hygromycin (400 μg/mL) (Roche, Cat#10843555001) for 7 days. The surviving cells were referred to as MEFs-dCas9 which contained the components of the CRISPR/SAM system^39^. The sgRNAs to activate *Hnrnpk* were designed using the online CRISPR design tool (http://crispr-era.stanford.edu/) and were cloned into lentiSAMv2 as previously described^39^. The correct insertion of the sgRNA sequences was confirmed by Sanger sequencing (performed at the Genomics Unit at CNIO). An empty plasmid lentiSAMv2 without sgRNAs was used as a nontargeting control. Lentiviral particle production was performed as previously described^40^. Lentiviruses were used to infect 3 × 10^5^ cells cultured in medium supplemented with 8 μg/mL polybrene and seeded in a 6-well plate. To mitigate possible biases due to off-target effects of the sgRNAs, *Hnrnpk*^SAM^ models were generated using two different sgRNAs per gene. The sequences of sgRNAs were as follows:

1. sgRNA1: Forward (5’-3’): CACCGCGCTGCTCACGTGTGCCGGG; Reverse (5’-3’): AAACCCCGGCACACGTGAGCAGCGC.
2. sgRNA2: Forward (5’-3’): CACCGCCGAGGGAGTTTGGCGCGAT; Reverse (5’-3’): AAACATCGCGCCAAACTCCCTCGGC.

The expression of *Hnrnpk* was quantified with qRT-PCR, and HNRNPK protein was measured by WB.

### RNA-seq analysis of hnRNP K overexpressing MEFs

Total RNA samples from empty vector and sgRNA2 MEFs were converted into sequencing libraries with the “NEBNext Ultra II Directional RNA Library Prep Kit for Illumina” (NEB #E7760), as recommended by the manufacturer. Briefly, the polyA+ fraction was purified and randomly fragmented, converted to double-stranded cDNA and processed through subsequent enzymatic treatments of end-repair, dA-tailing, and ligation to adapters. The adapter-ligated library was completed by PCR with Illumina PE primers. The resulting purified cDNA libraries were applied to an Illumina flow cell for cluster generation and sequenced on a NextSeq550 (Illumina) instrument in a single read 85 base format by following the manufacturer’s protocols. Adapters and remaining ribosomal sequences were removed with bbduk (http://sourceforge.net/projects/bbmap/). The resulting reads were analyzed with the nextpresso. pipeline as follows: sequencing quality was checked with FastQC v0.11.0 (https://www.bioinformatics.babraham.ac.uk/projects/fastqc/). Reads were aligned to the mouse genome (GRCm39) with TopHat2^41^ using Bowtie1^42^ and Samtools^43^, allowing 3 mismatches and 20 multihits. The Gencode vM29 gene annotation for GRCm39 was used. Read counts were obtained with HTSeq^44^. Differential expression and normalization were performed with DESeq2^45^, keeping only those genes where the normalized count value was higher than 10 in at least 30% of the samples. Finally, those genes that had an adjusted p-value below 0.05 FDRs were selected. GSEAPreranked was used to perform gene set enrichment analysis for the selected gene signatures on a preranked gene list, setting 1000 gene set permutations^18^. Only those gene sets with significant enrichment levels (FDR q-value < 0.25) were considered.

### Nucleolar stress assay

Briefly, MEFs (9,000 cells/well) were plated onto imaging 96-well plates (Perkin Elmer, #6055300). The following day, the cells were either preexposed or not to 5 nM ActD (Merck, #50-76-0) for 4 hr. Cells were fixed, blocked and stained according to the following immunofluorescence protocol.

### Global translation analysis by L-homopropargilglicine assay (HPG)

Briefly, MEFs (9,000 cells/well) were plated onto imaging 96-well plates. The following day, the cells were pulsed with HPG for 90 minutes, fixed with 4% PFA, and Click-iTTM chemistry was conducted following the manufacturers’ instructions for the Click-iTTM HPG Alexa FluorTM 488 Protein Synthesis Assay Kit (Thermo Fisher Scientific, C10428). Nuclei were stained with DAPI. After staining, the cells were detected by immunofluorescence.

### Polysome profile assay

Polysomal profiling analysis was performed by IMMAGINA BIOTECHNOLOGY. Briefly, cytoplasmic lysates were prepared by resuspending MEFs from 150 cm^2^ 70% confluent plates in 200 μl of IMMAGINA lysis buffer (IMMAGINA BIOTECHNOLOGY, Cat no. RL001-1). The cell suspension was passed through a G26 needle 10 times and cell debris and nuclei were pelleted by centrifugation at 20,000 x g for 5 min. Cleared supernatants were loaded on a linear 15%–50% sucrose gradient and ultracentrifuged in a SW41Ti rotor (Beckman) for 1 hr and 30 min at 180,000 g at 4°C in a Beckman Optima LE-80K Ultracentrifuge. After ultracentrifugation, gradients were fractionated in 1 mL volume fractions with continuous monitoring absorbance at 254 nm using an ISCO UA-6 UV detector.

### Immunofluorescence

Cells were fixed with paraformaldehyde (PFA) 4% in 1X PBS for 10 min at RT and permeabilized and blocked at the same time with a blocking buffer (BSA 0.5% and 0.1% Triton X-100 in 1X PBS) overnight at 4°C. After this process, the indicated concentration of primary antibodies in blocking buffer was applied: Hnrnpk (1:500, Santa Cruz, sc-28380), Ncl (1:400, Abcam #ab22758) and Fbl (1:250, Cell Signalling #2639). The following secondary antibodies were added overnight at 4°C in blocking buffer: Alexa Fluor™ 647 donkey anti-rabbit IgG (1:1000, Invitrogen, #A-32795), Alexa Fluor™ 488 donkey anti-mouse IgG (1:1000, Invitrogen, #A-21202). andstaining, cell nuclei were stained using DAPI (0.5 μg/mL, Invitrogen D1306) overnight at 4°C in blocking buffer.

Images were acquired within each cell. The total area of the nucleolus and number of dots were calculated by Ncl measurement. The nucleolar stress signal was analyzed by the increase in Ncl mean intensity in the nucleus of cells using DAPI as a counterstain. In these experiments, 20 pictures per well were taken using a 20X water objective. Image analysis was performed using a homemade pipeline identifying the different organelles according to the intensity of the signal in the nucleus. Images were taken using two High Content Screening System (HCS), HCS Opera™ LX and HCS Opera™Phenix® Plus and analyzed with their specific software: Acapella 2.6 (Perkin Elmer) and/or Harmony 5.1 (Perkin Elmer).

### Flow cytometry analysis

For cell cycle analysis, samples from MEF cultures (Hnrnpk^SAM^, Hnrnpk^Tg-cre^, wild type and empty vector) were collected and fixed with a glacial 70% ethanol solution overnight, followed by DAPI 40 ng/mL overnight incubation at 4°C. Then, the sample was passed through a strainer to ensure single cell suspension and analyzed using a FACS Canto II cytometer, recording 20.000 events per sample. For immunophenotyping experiments. We analyzed LSK (c-kit+/sca1+), LT-HSC (Cd34-/Sca-1+), ST-HSC (Cd34+/Sca-1+), myeloid-derived suppressor cells (MDSCs) (Gr-1+, Cd11b+), lymphoid (B220+), erythroid (ter-119+) and megakaryocytic (CD41+) compartments using a stem (HSC) and nonstem hematological cell panels (see Table 2). Samples were analyzed using an LSR Fortessa cytometer, recording more than 50.000 total events per sample. All data were analyzed with FCS Express 7 software (https://denovosoftware.com/)

**Table 2:** FCM antibodies

| Host | Name | Clone | Dye | Dilution | [ ] µg/mL | Reference |
| --- | --- | --- | --- | --- | --- | --- |
| **HSC panel** |  |  |  |  |  |  |
| Hamster | CD34 | HM34 | APC | 1:50 | 4 µg/mL | Biolegend 128612 |
| Rat | CD117 (c-Kit) | S18020A | PE | 1:100 | 2 µg/mL | Biolegend 161503 |
| Rat | Sca-1 (Ly-6A/E) | D7 | FITC | 1:100 | 5 µg/mL | Biolegend 108105 |
| Rat | CD127<br>(IL7Rα) | A7R34 | Brilliant Violet<br>421<sup>TM</sup> | 1:50 | 1 μg/mL | Biolegend<br>135023 |
| **Hematological cells panel** |  |  |  |  |  |  |
| Rat | Gr1 (Ly-6G/Ly-6C) | RB6-8C5 | APC | 1:400 | 0.5 μg/mL | Biolegend<br>108412 |
| Rat | CD11b<br>(MAC-1) | M1/70 | FITC | 1:400 | 1.25 μg/mL | Biolegend<br>101206 |
| Rat | B220<br>(CD45R) | RA3-6B2 | Brilliant Violet<br>510<sup>TM</sup> | 1:100 | 2 μg/mL | Biolegend<br>103248 |
| Rat | Ter119 | TER-119 | PerCP/Cyanine<br>5.5 | 1:40 | 5 μg/mL | Biolegend<br>116228 |
| Rat | CD41 | MWReg30 | PE/Cyanine7 | 1:1000 | 0.2 μg/mL | Biolegend<br>133916 |

### Senescence assays

MEFs were plated onto 6-well plates until 70-80% confluent. Senescence-associated (SA) β-galactosidase activity was determined by using the Senescence SA-β-Galactosidase Staining Kit (Cell Signaling, #9860), according to the manufacturer’s instructions. As senescent molecular markers such as p21 and p16 are expressed at low levels under steady-state conditions, we sublethally irradiated MEFs (*Hnrnpk*^SAM^) with 5 Gy and determined the levels of p21 and p16 protein with and without IR by western blotting.

### Western Blot

Western blotting was performed using protein lysates from MEFs and tissues such as the liver and spleen of wild type, Hnrnpk^Tg^ and Hnrnpk^-Cre^ mice. Tissues and cells were homogenized in RIPA lysis buffer (Millipore, #20-188) containing protease and phosphatase inhibitors (Complete Mini, Roche and PhosSTOP Roche, respectively). Soluble proteins were boiled in Laemmli buffer (Bio-Rad, Cat No. #1610747) with β-mercaptoethanol, resolved on a 4-20% gradient SDS-PAGE gel, and transferred to a PVDF membrane. Membranes were blocked in 5% nonfat milk for 1 hour at RT and then incubated with primary antibodies in bovine serum albumin (BSA) Tween-Tris buffered saline (T-TBS) 1X solution overnight at 4°C.

After primary antibody incubation, membranes were washed in 1X T-TBS and incubated in HRP-conjugated secondary antibody to be detected by the enhanced chemiluminescence SuperSignal ®West Femto Maximum Sensitivity Substrate (Thermo Scientific, Cat No. #34095). βActin and/or GAPDH were used as cellular loading controls.

The primary antibodies used were hnRNP K (Santa Cruz, D-6), p53 (Santa Cruz, DO-1), c-Myc (Abcam, ab32072), mTOR (Cell Signaling, #2983S), 4E-BP1 (Cell Signaling, #9644), p-4E-BP1(Cell Signaling, #2855), Ncl (Abcam, ab22758), p16(INK14a) (Thermo Fisher, MA5-17142), p21/CIP1 (Santa Cruz, F-5), β-actin (Cell Signaling, 4967S) and GAPDH (Cell Signaling, 2118S)

### qRT-PCR

Total RNA from frozen tissue and cell cultures was extracted using the RNeasy kit (QIAGEN, Cat No. #74134) following the manufacturer’s instructions. cDNA synthesis was performed using the iSCRIPT cDNA synthesis kit (BIO-RAD) according to the manufacturer’s protocols. Quantitative real-time PCR was performed with the QuantStudio 6 Flex (Applied Biosystems, Life Technologies) using SYBR qPCR master mix (Promega, Cat No. #4367659). All values were obtained in triplicate. Primers for mouse samples are listed in Table 3.

**Table 3:**
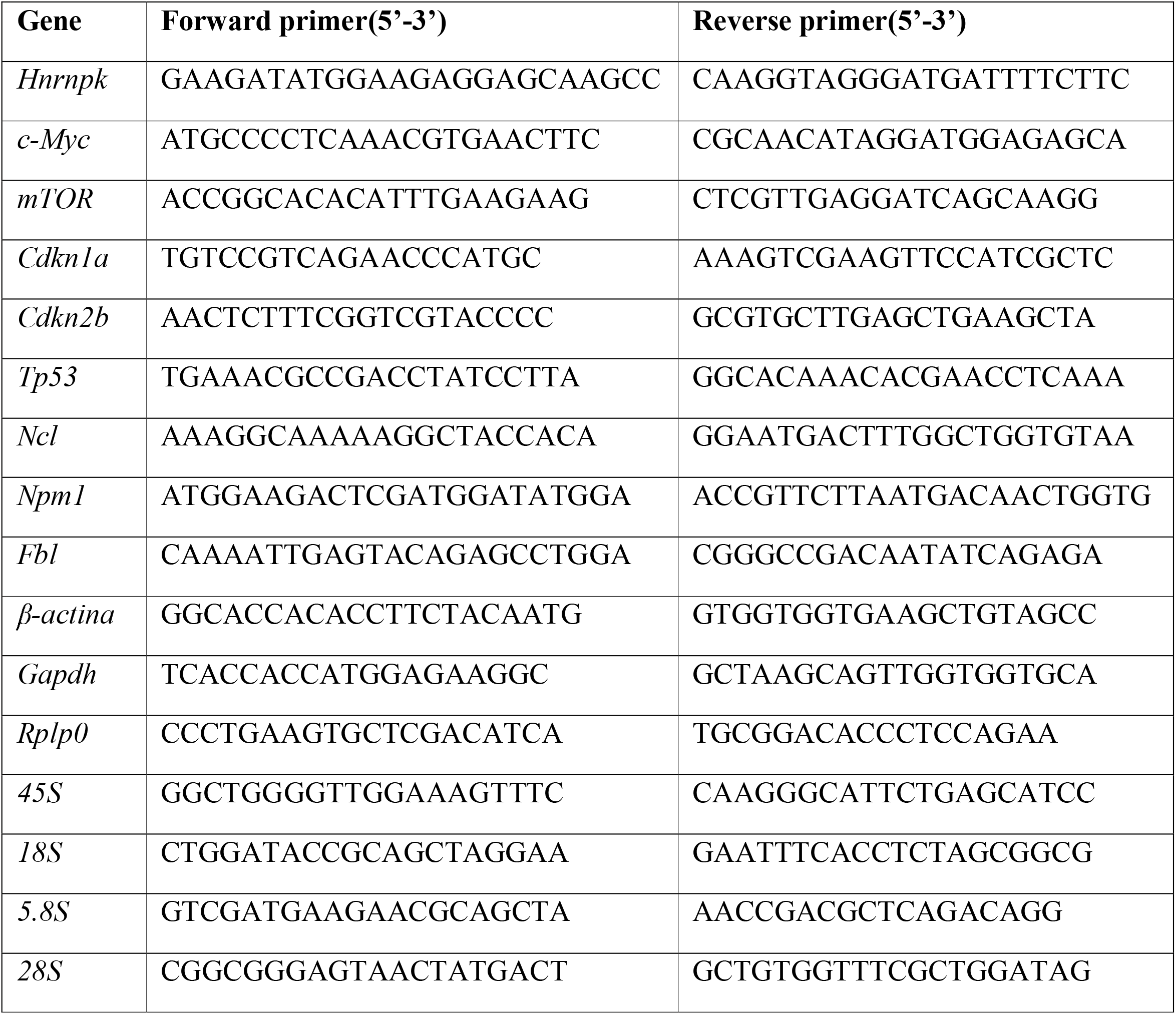
qPCR primers

*Gapdh* and/or β*-actin* served as housekeeping controls. All experiments were performed in technical triplicates. Changes in expression were compared using the Pfaffl method^46^ by comparing expression changes between target genes and the housekeeping control.

### Quantification and statistical analysis

Statistical analyses when comparing two groups were performed using unpaired t-tests and Mann-Whitney tests, with the exception of populations with normal distribution (p-values less than 0.05 in the normality KS test). For dichotomic discrete variables, chi-square tests were performed, and p-values less than 0.05 were considered statistically significant. For multiple group analysis a two-way ANOVA Sidak’s multiple comparisons test was performed. To test differences in survival curves the Kaplan-Meier test was used. Differences between survival distributions were analyzed using the log-rank test. Hazard ratios and confidence intervals were obtained by Mantel-Haenszel analysis. Statistical computations were performed using GraphPad Prism 7.0.

## Acknowledgments

The authors would like to thank CNIO whole animal facility core unit staff and especially members Isabel Blanco, Gema Luque, Gema Iglesias and Yolanda Cecilia for their support in animal care; CNIO Histopathology Unit personnel for their support in histopathology and immunohistochemistry analysis. Genomics Unit personnel for genotyping and sequencing; CNIO flow cytometry Unit head, Lola Martinez, and staff for their FACS support; CNIO Genomic Instability Group members Matilde Murga, and Oscar Fernandez-Capetillo for their helpful advice; and nurse Carmen Delgado for her histopathology advice.

